# Assessing the Efficacy of Computational Workshops and Participatory Live Coding in Evolutionary Biology

**DOI:** 10.64898/2026.05.04.722624

**Authors:** Sarah Swiston, Lisa Kuehne, Rick Moore, Michael Landis

## Abstract

Computational workshops are common in evolutionary biology and are used to share discipline-specific tools and skills with researchers. Despite the perceived importance of these workshops, there is no common set of criteria for workshop success, and there are few peer-reviewed studies investigating the efficacy of workshops or assessing the value of particular instructional techniques in this context. Here, we focused on one key element of a successful workshop: its ability to increase participants’ motivation to use the methods and tools presented during the workshop. We analyzed the goals, perceptions, and future plans of research practitioners engaging in a workshop on phylogenetic methods of historical biogeography using pre- and post-workshop surveys. Overall, the workshop was successful at motivating participants, and survey responses provided insights into participants’ perceptions of different activities, including “participatory live coding”. Apart from this case study, we aim to highlight the importance of developing a common set of workshop goals in collaboration with other workshop stakeholders and the need for specialized, validated tools for assessing the efficacy of computational workshops for researchers.

## Introduction

In the field of evolutionary biology, computational skills are often required to complete fundamental aspects of the research process such as data analysis, visualization, and simulation (National Research Council, 2011; Wright et al., 2019; Dabholkar et al., 2021; McDonald et al., 2022). Many undergraduate biology programs require minimal (if any) coursework related to statistics, computation, programming, or mathematical modelling (Wilson Sayres et al., 2018). Even when these general skills are intentionally sought by undergraduates, the amount of field-specific knowledge necessary to engage in research is often beyond the scope of undergraduate training. Thus, many early-career researchers, including graduate students and postdoctoral researchers, rely on specialized instruction from a variety of sources. Alongside upper-level coursework, direct mentorship, literature review, and tutorials, computational workshops are thought to play an important role in teaching early-career researchers how to use discipline-specific computational methods and tools.

Facilitators, participants, and funding organizations clearly believe that workshops provide value. They commit a substantial amount of time and resources to workshops, and would likely be interested to know whether those investments are worthwhile. However, workshop stakeholders do not always articulate the broader purposes of workshops, and to our knowledge, there is no shared set of goals that make computational workshops for researchers “successful”. Without common standards or tools for assessing whether a particular workshop is successful, it is challenging to compare amongst workshops or assess the efficacy of different instructional techniques. Our view is that that workshops for researchers have different goals than classroom instruction, and the conventional definition of success based on students’ performance on a set of learning objectives cannot be directly applied. Workshop instructors generally understand this; they typically avoid exams or assignments to assess performance, instead favoring surveys to learn about participants’ expectations, experiences, and understanding. These surveys are very useful for iteratively improving instruction, but are not designed to assess a common measure of workshop success in a standardized way. **In this study, we focus on the goal of *increasing participant motivation*, and provide an example for how to measure it in the context of a computational workshop for researchers**. Ultimately, a common set of goals should be developed by workshop stakeholders through focus groups and surveys, and assessment tools should be appropriately validated.

Drawing on our own experiences with computational workshops, we identified four primary aims of workshops: 1) to orient members of the field to some of the research methods and tools available to them, and what those methods can and cannot do, 2) to help them implement those methods and tools if the researcher determines that they are appropriate for their own research questions, 3) to give them new ideas of questions they had not considered which could be addressed using existing methods and tools, and 4) to empower them to apply methods and tools which the workshop organizers consider valuable. While there are other goals of any good workshop, such as improving diversity and equity and creating new networking opportunities, one important outcome of a successful workshop might be whether it results in researchers generating high-quality research that implements the tools or methods they gained during the workshop. Advancing the field through the production of new research is important to all major stakeholders, including the instructors who facilitate workshops, the participants who attend them, and (if applicable) external funding sources.

Unfortunately, the link between a workshop or intervention and the production of high-quality research is difficult to examine. When researchers are designing a project, they must determine which methods are appropriate to answer their specific questions and how to appropriately apply those methods in the context of their own work. Attending workshops and consulting with instructors and fellow participants can certainly help researchers determine whether and how to use methods and tools, but the fate of a research project is not immediately measurable. Investigators continuously evaluate their own work and encounter many choices and barriers during the course of a research project. Their decision to pursue a particular research direction may be affected by external challenges, such as resource limitations. Once a project has been completed, its assessment is then the responsibility of peer reviewers during and after publication. It may be useful to track the number of studies published by workshop participants involving methods taught during a workshop, the quality of the journals where those studies are published, and the number of citations generated. However, publication success is the result of many factors which are difficult to disentangle, such as the differing visions of journal editors, opinions of reviewers, and biases toward certain types of research or researchers (Ferber & Brün, 2011; Gomez et al., 2022). These elements pose a substantial challenge to a workshop instructor who wants to use participants’ publications to assess the effectiveness of a workshop and its activities. Rather than publication success, we suggest that participant motivation is a more direct reflection of a workshop’s ability to promote high-quality research implementing particular methods and tools. **A successful workshop should increase participants’ motivation to use the methods and tools presented during the workshop**.

If participant motivation is a key element of a successful workshop, our assessments should reflect this. While motivation can be tied to ability, one would not expect immediately measurable workshop learning objectives, such as the learner’s ability to complete coding tasks or answer questions about the details of a method, to predict motivation. A researcher who sees the value in a particular method and feels confident about their capabilities will typically do further reading about the method they intend to use, and might troubleshoot until their implementation is functional, even if it is difficult for them. A researcher who does not see value in a method, or who does not feel confident about their capabilities, might not even try to implement a tool or method at all. Here, we implicate the situated expectancy-value theoretical model, which postulates that achievement-related decisions are influenced in complex ways by both expectancies and values (Eccles & Wigfield, 2023). Expectancies are defined as an individual’s beliefs about how well they will do on a particular task; they are driven by many factors, including an individual’s self-concept of their own abilities and their perception of task difficulty. Values, on the other hand, are influenced by an individual’s perceptions of four primary components: task importance, enjoyability, usefulness, and cost. Under this framework, we expect that a workshop participant’s motivation to use tools or methods from the workshop should be influenced by their perceptions of several key factors, including their ability to use the methods, the difficulty of using the methods, and usefulness or applicability of the methods for their own research.

Several studies have examined the efficacy of introductory-level computational workshops in biology through surveys about participant perceptions. Oesper & Vostinar (2020) found that undergraduate students (N=22) participating in a computational workshop consisting of short modules from several biological fields considered themselves more likely to pursue future opportunities related to computational biology. Zuvanov et al. (2021) surveyed undergraduate and graduate students (N=37) attending an introductory Python workshop in Brazil. Participants reported that writing their own code was helpful, and the workshop was applicable to their research. Timms & Guyon (2023) observed similar responses amongst graduate students (N=25) attending another introductory Python workshop. These studies point toward the usefulness of introductory-level computational workshops. However, the factors that motivate an undergraduate student to enter a new field, or those that prompt a student to try coding, will inevitably differ from the factors governing experienced researchers’ decisions to implement cutting-edge methods.

It is also common to administer surveys to participants attending advanced-level workshops. For example, the Woods Hole Workshop on Molecular Evolution (https://molevolworkshop.github.io/), one of the largest and longest-running workshops in the field, frequently surveys its participants, as do the RevBayes workshops (Barido-Sottani et al., 2022; Workshop on Molecular Evolution, 2025). These surveys are useful for modifying future iterations of a workshop, but are not publishable, IRB-approved research studies designed for generalizability. Studies assessing the efficacy of advanced-level workshops are rare. Magnano et al. (2022) focused on a machine learning workshop for biologists, surveying participants (N=47) before and after the workshop to assess how their confidence and plans changed.

Participants indicated that they felt more knowledgeable and confident after the workshop, and that they were more likely to identify and solve problems related to machine learning. The study also identified common challenges and provided recommendations to workshop administrators, demonstrating the value of conducting research in this area.

To address the lack of research on the efficacy of advanced-level computational workshops, we used pre- and post-workshop surveys to investigate participants’ perceptions and plans regarding a computational workshop for practicing researchers in evolutionary biology, with the goal of identifying whether this workshop was effective according to our motivation-based criteria.

Generating information about the value of specific activities is also important for informing future workshop design. Therefore, it is important to ask learners about their perceptions of individual activities and whether those activities influenced their research plans. Most workshops contain a variety of activity types, including lectures, problem-based exercises, discussion, and live coding demonstrations. One frequently employed activity is “participatory live coding”.

Participatory live coding falls into the broader category of “live coding”, a set of techniques where the instructor types code actively during instruction. This is in contrast to “static coding”, where code is presented in finished form (the lecturer is not typing the code live). During live coding, an instructor types out lines of code on a computer that is visible to learners (usually projected on a screen, or in the case of online instruction, via screen-sharing). It is important that the instructor actively types the code for two main reasons: it slows the instructor down, and it allows for “organic” mistakes to occur, giving learners a good example of “debugging” (Nederbragt et al., 2020). This active coding is accompanied by verbal instructions, where the instructor describes what they are doing as they type. This is important because there is strong evidence that receiving information in multiple ways is beneficial for learning (Clark & Paivio, 1991). The instructor gives their reasoning behind each coding step, interspersed with more general reasoning about the process as a whole. There is some research demonstrating the value of non-participatory live coding in computer science. Studies find that live coding produced better learning outcomes than both traditional lecturing and “static coding” (Rubin, 2013; Raj et al., 2018, 2020).

“Participatory live coding” is a specific type of live coding where participants “code along” with the instructor, typing the same code into their own computer and running it themselves. They may also take notes or write code comments with explanations. This technique often involves taking questions from participants who are struggling to get their code to run, and extra instructors may be present for this purpose. The participatory live coding can also be interspersed with small exercises that learners are expected to complete on their own, followed by a group discussion or instructor-led lesson about one way the problem can be solved. This strategy combines the benefits of challenge-based learning with the benefits of guided instruction (Kirschner et al., 2006; Connell et al., 2016; Serrano et al., 2018).

Participatory live coding is a favored tool amongst certain workshop designers who wish to enable participants to apply particular computational tools to their datasets in the future. The Carpentries, a global project consisting of three instructional organizations (Data Carpentry, Library Carpentry, and Software Carpentry), uses participatory live coding as a primary tool for its computational workshops (Wilson, 2014, 2019). Carpentries workshops for researchers are focused on teaching participants how to use software tools. They are different from the advanced-level computational workshops which expose researchers to discipline-specific methods with the goal of promoting research in the field. However, we might expect that researchers would experience the participatory live coding activity similarly. Carpentries administers surveys at their workshops, and their findings suggest that the workshops are perceived positively (Jordan et al., 2018). Unfortunately, these surveys do not address participatory live coding specifically.

There are few studies which specifically examine “participatory live coding”, and these studies are about classroom instruction, not workshops (Brown 2022). Because classroom instruction typically centers learning outcomes, the current research on live coding primarily assesses those outcomes. The studies may survey students, but those surveys are used to generate information about learning outcomes; the studies do not focus on students’ perceptions of their own learning in terms of empowerment or plans to implement learned skills following live coding activities. Understanding the impact that workshop activities have on researcher motivations requires additional research.

In this study, we examined the usefulness of computational workshops in the field of evolutionary biology by surveying participants attending the Phylogenetic Biogeography Workshop, a workshop about phylogenetic methods of historical biogeography (the study of the distribution of species in space over time). The survey investigated participants’ perceptions of their own learning and plans for implementing skills obtained during the workshop as a whole, and also addressed the “participatory live coding” activity specifically. We assessed whether workshop participants considered themselves more likely to implement phylogenetic models of historical biogeography and the corresponding software tools after engaging in the workshop, and whether the participatory live coding activities, in particular, were useful and/or motivating. We found that participants felt motivated to implement relevant models in the future, and their confidence improved in key areas. The participants also indicated a pervasive interest in learning about software tools, which was not a main advertised goal of the workshop. Despite this, the findings regarding participatory live coding were complex. The activity was perceived as helpful overall. However, when participants were asked how much “easier” it would be to implement the biogeographic models or how much “more likely” they were to do so, the null hypotheses of equal positive and non-positive responses could not be rejected. Participants generally preferred traditional lectures over coding activities, and some expressed frustrations with participatory live coding. These results highlight the importance of surveying participants to learn about how their motivation to use particular tools and methods is impacted by workshops and the instructional techniques therein. They also call for the development of standardized, validated tools for pedagogical research in the setting of computational workshops for researchers.

## Methods

As part of National Science Foundation (NSF) grant DEB 2040347, three computational workshops about phylogenetic methods of historical biogeography (entitled “Phylogenetic Biogeography Workshops”) were administered. The workshops highlighted several mathematical models describing how species evolve and move through space, including birth-death models and continuous-time Markov chains, along with several biogeographic models with acronyms as names (DEC, GeoSSE, FIG, and MultiFIG). Lessons drew upon numerous advanced topics from fields such as molecular phylogenetics, diversification modeling, historical biogeography, Bayesian inference, stochastic processes, and programming. We used a backward design approach and generated a series of lessons that built toward implementing a complex FIG or TimeFIG analysis (Wiggins & McTighe, 1998). The workshops covered the same topics in roughly the same manner, with the primary differences being time and location: a 3-day workshop was given at Washington University in Saint Louis (WashU, June 4–6, 2024), and 1-day workshops were given at the Evolution 2025 conference (June 20, 2025) and the Botany 2025 conference (July 27, 2025).

The workshops were offered free of charge, although the conference-based workshops required participants to be registered for the conference, and travel expenses (excepting hotel accommodations for the WashU workshop) were not covered. The application process for the 3-day workshop required potential participants to describe their research goals and rate their experience with certain topics so that the facilitators could choose a cohort of applicants with similar abilities and sufficient relevant experience to benefit from the workshop. In particular, attendees were required to have some prior knowledge of phylogenetics and programming, and to be interested in biogeography. After determining which applicants qualified, participants were randomly selected. The application materials were not used to investigate the efficacy of the workshop because participants may present their research plans and experiences differently during the application process than during a survey without any stakes. Additionally, we did not want applicants to worry that study participation would influence workshop admission, so the research project was not introduced until after workshop participants were selected. Because the 1-day workshops took place at conferences, admission was handled through conference registration. Therefore, participants attending the 1-day workshops were not required to submit applications, and all registrations were accepted until the available seats were filled.

The workshops were advertised through multiple channels, including social media, institutional websites, word of mouth, and conference registration forms. The advertisements indicated that the workshop was about phylogenetic biogeography, and all participants should have some background in phylogenetics and programming. It was also stated that, while the workshop would be conducted in RevBayes (a programming language, Höhna et al., 2016), participants were not required to have any prior experience with this tool. The workshop was not advertised as a “RevBayes workshop” (Barido-Sottani et al., 2022).

The workshop content was designed for researchers with some experience in phylogenetics and programming, although levels of knowledge varied. Advanced undergraduates, graduate students, postdocs, and early career faculty were welcome to apply; our pre-workshop survey respondents across all three workshops consisted of 1 undergraduate student, 1 post-baccalaureate researcher, 32 graduate students, 8 postdoctoral researchers, 3 faculty members, and 2 non-faculty professionals. Participants were asked to identify their career stage during the pre-workshop survey in case participants’ goals, experiences, workshop benefits, and future plans differed depending on their career stage. As an example, a graduate student may be more likely to change research directions based on a positive workshop experience. Alternatively, a postdoctoral researcher may be able to complete a full analysis by the end of the workshop, making them more likely to publish related work in the immediate future. However, our final sample size did not allow us to distinguish these effects.

Before each workshop, participants were provided with the required software tools and a series of written online tutorials that would prepare them to engage with the workshop content. The completion of these tutorials was not assessed. Virtual office hours were also held to assist with software installation, which were attended by a small number of participants. The workshop itself was designed to equip participants with the necessary background knowledge and tools to conduct phylogenetic analyses of historical biogeography. The material was presented through a variety of activities, including lectures, discussions, coding tasks, and participatory live coding.

To examine the efficacy of the workshop and participatory live coding activity, researchers administered two surveys: a pre-workshop survey **(Supp. Mat. S1)**, and a post-workshop survey **(Supp. Mat. S2)**. The surveys were emailed electronically via the WashU instance of REDCap (Research Electronic Data Capture) to all participants prior to and immediately following each workshop (Harris et al. 2009; Harris et al. 2019). The surveys were accompanied by information about the research project **(Supp. Mat. S3)**. At the beginning of the workshop, a verbal announcement was made to participants acknowledging the optional surveys, providing the general purpose of the surveys and their use in a pedagogical research project, and encouraging participants who had not filled out the pre-workshop survey to do so **(Supp. Mat. S4)**. The pre-workshop survey was sent three days before the workshop and remained open until the end of the first “coffee hour” or “introductions” section of the workshop. Another announcement was made after the workshop, indicating that the post-workshop survey had been sent electronically **(Supp. Mat. S4)**. The post-workshop survey was sent immediately following the conclusion of workshop activities. Participants were given ample time to complete the post-workshop survey, either immediately following the workshop (in the computer lab where the workshop took place), or any time within the next week. Consent was acquired prior to the administration of each survey. Survey participants had the option to skip any questions they did not wish to answer, and participants were asked to directly consent to the use of verbatim responses in publication. Data was anonymized before analysis, with pre-and post-workshop surveys linked only via anonymous identifiers. This work was classified as exempt from IRB review by the Washington University in Saint Louis IRB (ID# 202405112).

The first workshop had 21 registrants (of 63 applicants), the second workshop had 24 registrants, and the third workshop had 26 registrants, for a total of 71. Some of these registered individuals were unable to attend the workshop; any pre-workshop survey responses received from non-attendees were excluded. N/A options were offered on survey questions regarding individual workshop activities that participants may have missed. In total, 47 participants completed the pre-workshop survey, and 31 participants completed the post-workshop survey, with 4 additional partial responses. In total, 49 participants were involved in one or both surveys. While participants were able to freely opt out of completing the surveys without consequences, response rates for both the pre- and post-workshop surveys were greater than 50%. The surveys included both multiple-choice (or rating) questions and short-response questions. Survey questions focused on three primary areas.

### Workshop Goals

During the pre-workshop survey, participants were asked to identify their initial goals for the workshop, choosing all that applied from a list of suggested goals with the option to write in their own response. They were also asked a multiple-choice question about whether they planned to use phylogenetic biogeography in their research and, if so, when they planned to start. This was followed by an open question about how participants intended to apply the methods to their own research. Theses question investigated the perceived purpose of the workshop, from the perspective of the participants. Participants were also given a set of possible research challenges related to implementing complex models, and were asked to rate how often they either stopped due to those challenges or continued despite them. This question was designed to assess whether a motivated researcher is likely to work through challenges; if so, this would highlight the importance of motivation over content knowledge.

During the post-workshop survey, participants were asked in an open question whether the workshop met their expectations and why. These responses were coded into five categories: “exceeded expectations”, “met expectations”, “partly met expectations”, “did not meet expectations”, and “other” responses that did not address the question as intended. Participants also rated how likely they were to use any skills or tools from the workshop. Additionally, participants rated whether they were more or less likely to engage with phylogenetic biogeography after attending the workshop, compared to how they felt before, alongside an open question about their reasoning. These responses allowed us to assess whether the workshop was perceived as useful by participants, and whether it successfully motivated participants to use phylogenetic models of historical biogeography.

### Workshop Content

In the pre-workshop survey, participants were asked to rate their experience with content areas, including concepts (e.g. biogeographic models) and tools (e.g. RevBayes). In the post-workshop survey, participants were asked to rate how their level of confidence regarding these topics changed after the workshop. They were also asked open questions about their future plans for implementing tools and skills from the workshop: which ones the participants felt they could implement easily, and which ones they planned to use. These questions broke down the findings by topic to determine whether the key topics identified by instructors and participants were adequately addressed and inform future workshop development.

### Individual Activities

In the post-workshop survey, participants were asked to rate the effectiveness of individual workshop activities, including participatory live coding. Additionally, a description of “participatory live coding” was provided, and the participants were asked about the effectiveness of this specific activity through two questions: one asked them to rate whether the activity would make implementing models easier or less easy in the future, and the other asked them to rate whether they were more or less likely to use models after doing the activity. These questions were designed to assess the effectiveness of various instructional techniques for motivating participants, highlighting participatory live coding in particular.Participants were also asked to list any challenges they encountered during the workshop, providing more detailed information about their perceptions of workshop activities and aiding in the design of future workshops.

Data analysis was performed in R (R Core Team, 2025). Statistical analyses were not possible for certain research questions because of the small sample size (e.g. differences between different demographics of research participants). When necessary, response categories were grouped to facilitate statistical testing. All exact Wilcoxon signed rank tests were single-tailed using paired samples. All exact binomial tests were single-tailed with a null hypothesis of equal positive and non-positive results.

## Results

In the pre-workshop survey, participants indicated moderate experience with workshop topics (Fig. 1). Across topics other than the modeling language RevBayes, most participants were either “somewhat familiar” or “competent” (∼83%, p<0.001, exact binomial test), while few rated themselves as “highly competent” or “expert” (∼10%, p<0.001, exact binomial test). The most unfamiliar topic was RevBayes, the programming language used for the workshop (mean response ∼1.68/5, p<0.001, exact Wilcoxon signed rank test comparing RevBayes to the next most unfamiliar topic, biogeographic models). While teaching RevBayes was not a primary goal in the design of the workshop, when asked to select their goals in applying for the workshop from a list of possible goals, most respondents indicated that learning more about software tools was a goal for them (∼79%, p<0.001, exact binomial test). Most participants also indicated that learning about models (∼85%, p<0.001, exact binomial test) and learning about scientific questions (∼83%, p<0.001; exact binomial test) were amongst their goals.

**Figure 1.**
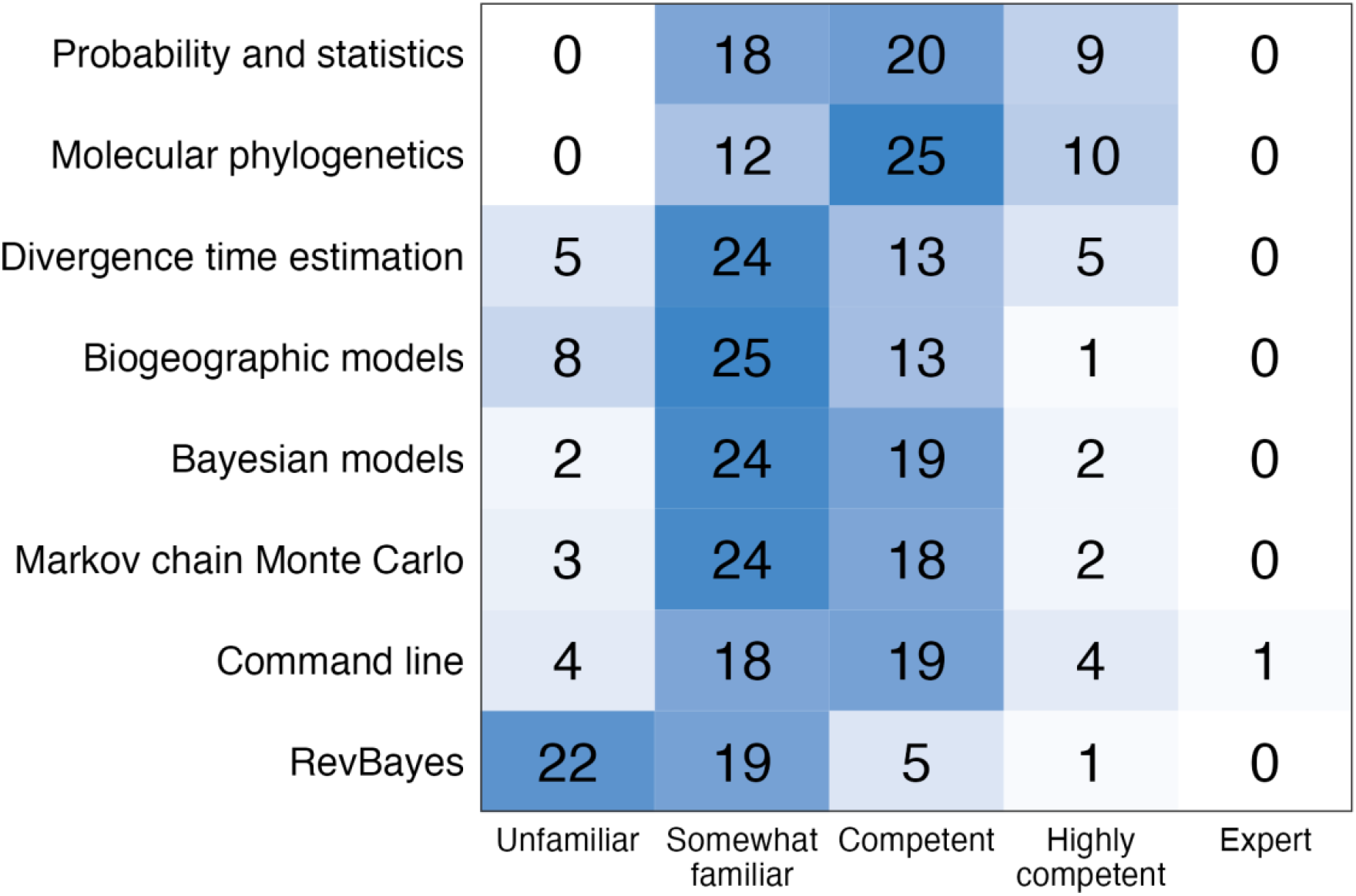
Participants’ prior experience levels across workshop topics. Most participants rated themselves as either “somewhat familiar” or “competent” across topics, excluding RevBayes (p<0.001, exact binomial test). The most unfamiliar topic was RevBayes, the programming language used in the workshop (p<0.001, exact Wilcoxon signed rank test).

Interestingly, only 18 of 47 pre-workshop respondents (∼38%) chose “decide whether to do future research using phylogenetic methods of historical biogeography” amongst their goals. Most participants indicated that they already planned to use these models prior to the workshop (∼74%, p=0.001, exact binomial test). Still, the results of the post-workshop survey showed that participants considered themselves more likely to engage with these models after having taken the workshop, compared to how they felt before (mean response ∼4.19/5, p<0.001, exact binomial test). One participant reported, “Now these models don’t feel like a dark, unknown place. I am aware that I still need to learn more in terms of the computational tools and mathematical characteristics of the models to fully understand them, but it was a really nice first approach for them not to feel so scary as before.” Another answered, “I had an understanding of biogeography and phylogenetic modeling but wasn’t aware of the sheer power and complexity of questions that could be asked!” Participants also highlighted the importance of being exposed to software tools. One said, “I had never used RevBayes, and the workshop was effective in showing how to install and use it.” Another responded, “I know more about it than I did before and have seen that it’s not that complicated to implement with PhyloDocker. Before, the software requirements were pretty intimidating.”

Participants also indicated higher confidence in the area of biogeographic models, which was a primary goal in the design of the workshop (mean response ∼4.27/5, p<0.001, exact binomial test) (Fig. 2). Most participants indicated that they were likely to use skills or tools acquired during the workshop in their future research (∼87%, p<0.001, exact binomial test). Participants who indicated that they would use skills or tools were asked a free-response follow-up question about *which* tools they planned to use, and 23 of 28 respondents referred to specific models that were taught during the workshop. Many participants believed they could directly apply the models from the workshop in their own research. For example, one participant said, “I definitely could use the MultiFIG model!” Another said, “I think I’d be able to run an analysis very similar to the ones we used (sort of ‘out of the box’) without too much difficulty… probably.”

**Figure 2.**
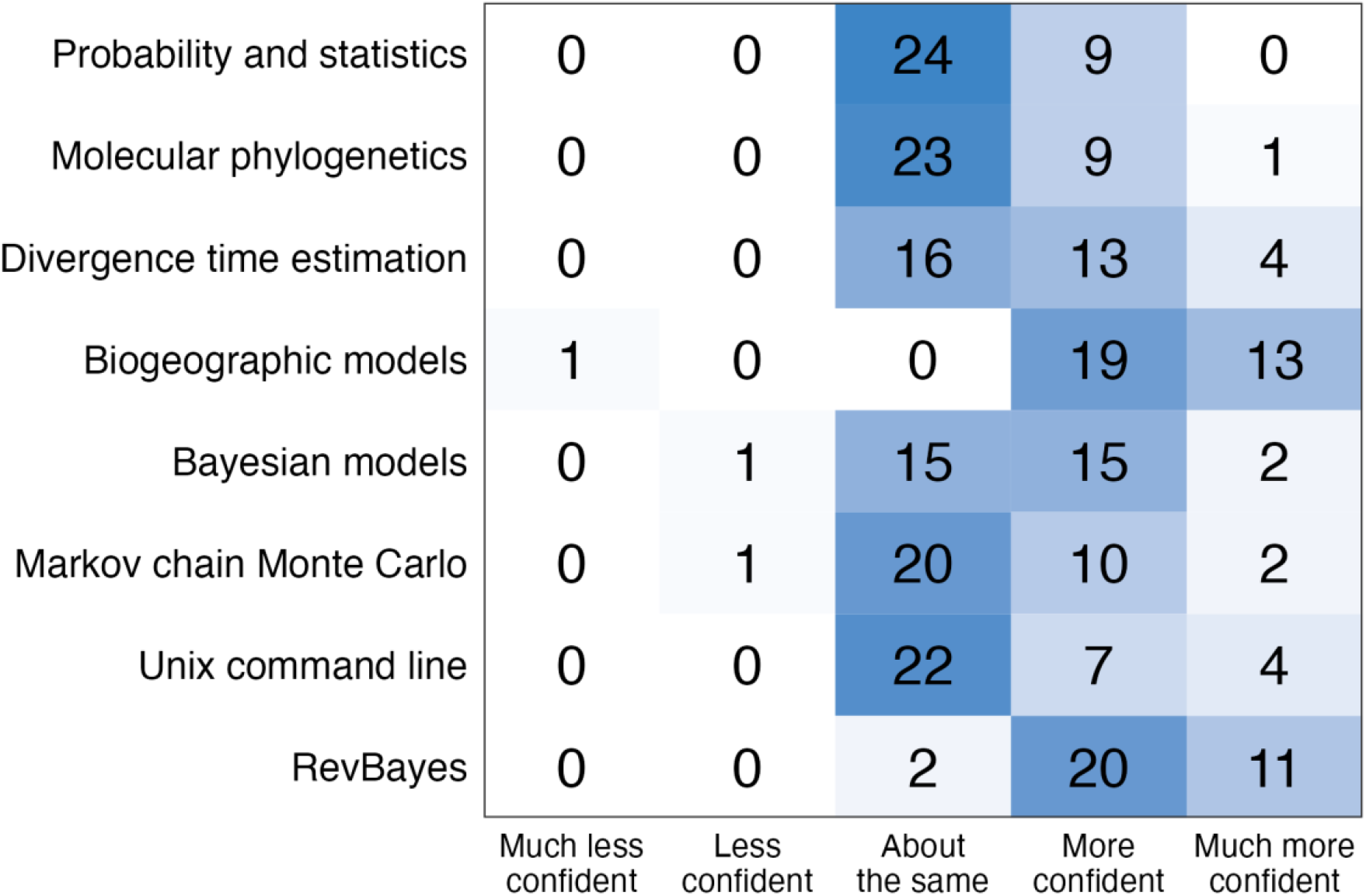
Participants’ changes in confidence across workshop topics. Participants reported increases in confidence regarding several topics, including biogeographic models (p<0.001, exact binomial test) and RevBayes (p<0.001, exact binomial test). Learning about biogeographic models was a primary goal for both instructors and participants. Learning RevBayes was a goal of most participants, but was not a major goal advertised by instructors.

Participants also felt that their own goals for the workshop were met (∼78%, p=0.001, exact binomial test). When asked why the workshop did or did not meet expectations, one participant said, “When I attended the workshop I was expecting to get a better understanding on how these analyses work in order to apply them on my own research. Now, I keep asking myself questions about the best way of gathering data to run the analyses and how to adapt the example we worked on to my own research system. This is only possible because I got a good basic sense on what the analyses are doing.” Another responded, “This workshop exceeded my expectations, I wasn’t aware that we would be learning about such new programs that would be able to really change the questions I want to ask about my systems and provide more precision than the biogeography analyses I’ve learned about previously. I also felt that the explanation of how to set up scripts in RevBayes and make all of that work was really well done especially as someone who had only a passing understanding of Bayesian statistics.” For the two topics with the lowest prior experience (RevBayes and biogeographic models), participants indicated that their confidence improved after the workshop (mean responses of ∼4.27/5, p<0.001 and ∼4.30/5, p<0.001; exact binomial test) (Fig. 2). Learning about software tools and models were the two most prevalent participant goals, and participants’ confidence increased in both of these areas.

These results point to the overall success of the workshop in encouraging participants to engage with the models taught during the workshop. The value of participants’ improved confidence is supported by their responses regarding persistence in the face of research challenges. Participants indicated that they were significantly more likely to continue research despite a variety of challenges than to stop due to those challenges (mean responses ∼3.23/5 and ∼2.48/5 respectively, p<0.001, exact Wilcoxon signed rank test) (Fig. 3). Therefore, the research team expects participants’ self-reported confidence and plans to be an indicator of their potential to do future research in the field, irrespective of the degree to which the material was “mastered” during the workshop.

**Figure 3.**
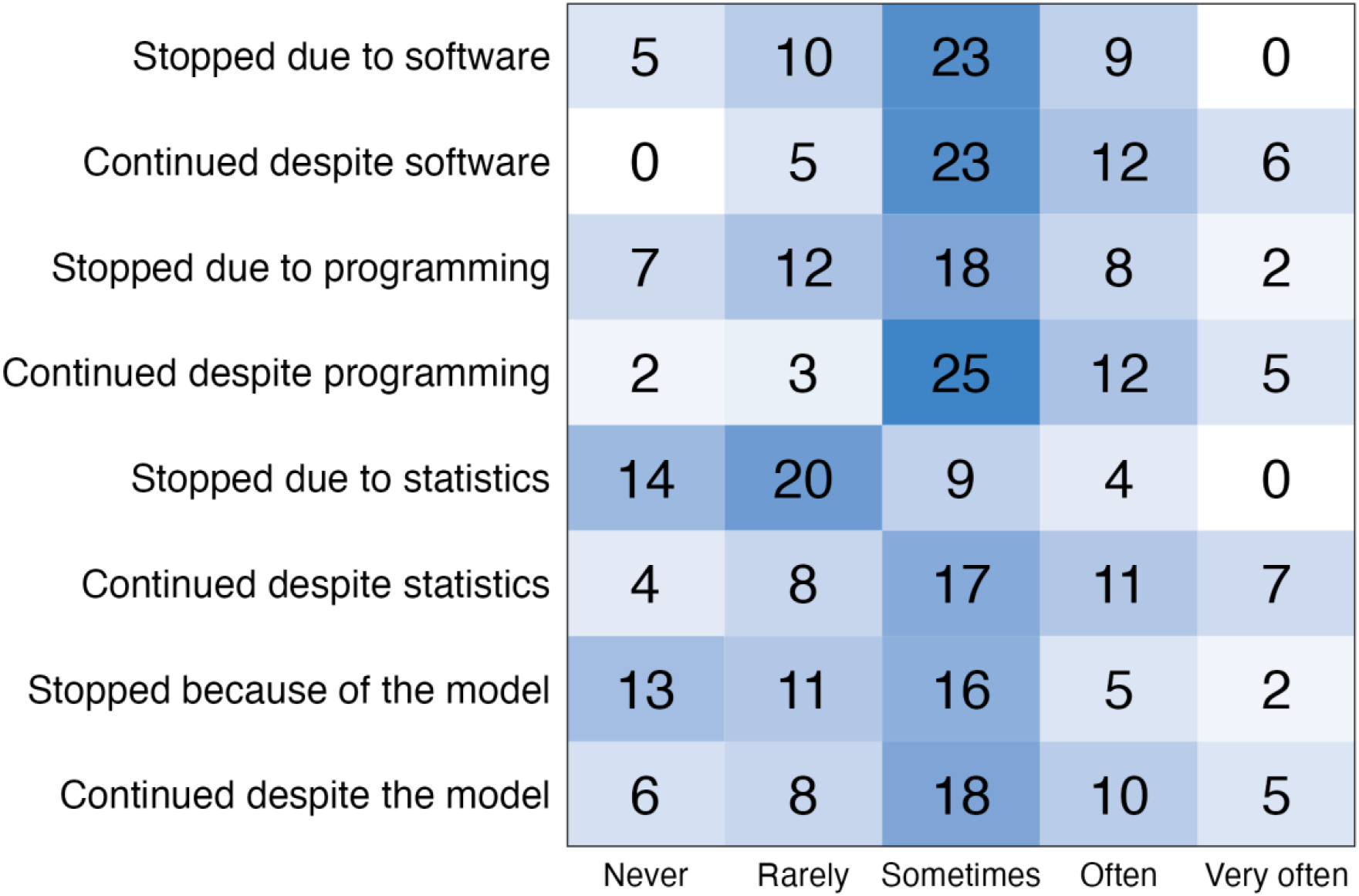
Participants’ responses to research challenges. Participants reported that they were significantly more likely to continue despite research challenges than to stop due to those challenges (p<0.001, exact Wilcoxon signed rank test).

When asked about the helpfulness of specific activities, participants rated lecture-style presentations more highly than other activities (mean responses ∼4.65/5 and ∼4.25/5 respectively, p=0.003, exact Wilcoxon signed rank test). Participatory live coding was considered helpful (mean response ∼4.06/5, p=0.015, exact binomial test), although one participant considered it unhelpful and a second participant considered it very unhelpful (Fig. 4). When asked specifically about participatory live coding, some participants said participatory live coding would make it easier to implement the models in the future (∼58%). However, one participant said the activity would make it more difficult to implement the models, and another said the activity would make it much more difficult. The null hypothesis of equal positive and non-positive outcomes could not be rejected (mean response ∼3.48/5, p=0.237, exact binomial test). Similarly, some participants said participatory live coding made them more likely or much more likely to engage with the models in the future (∼61%). However, one participant said the activity made them much less likely to engage with the models, and the null hypothesis of equal positive and non-positive outcomes could not be rejected (mean response ∼3.68/5, p=0.141, exact binomial test) (Fig. 5).

**Figure 4.**
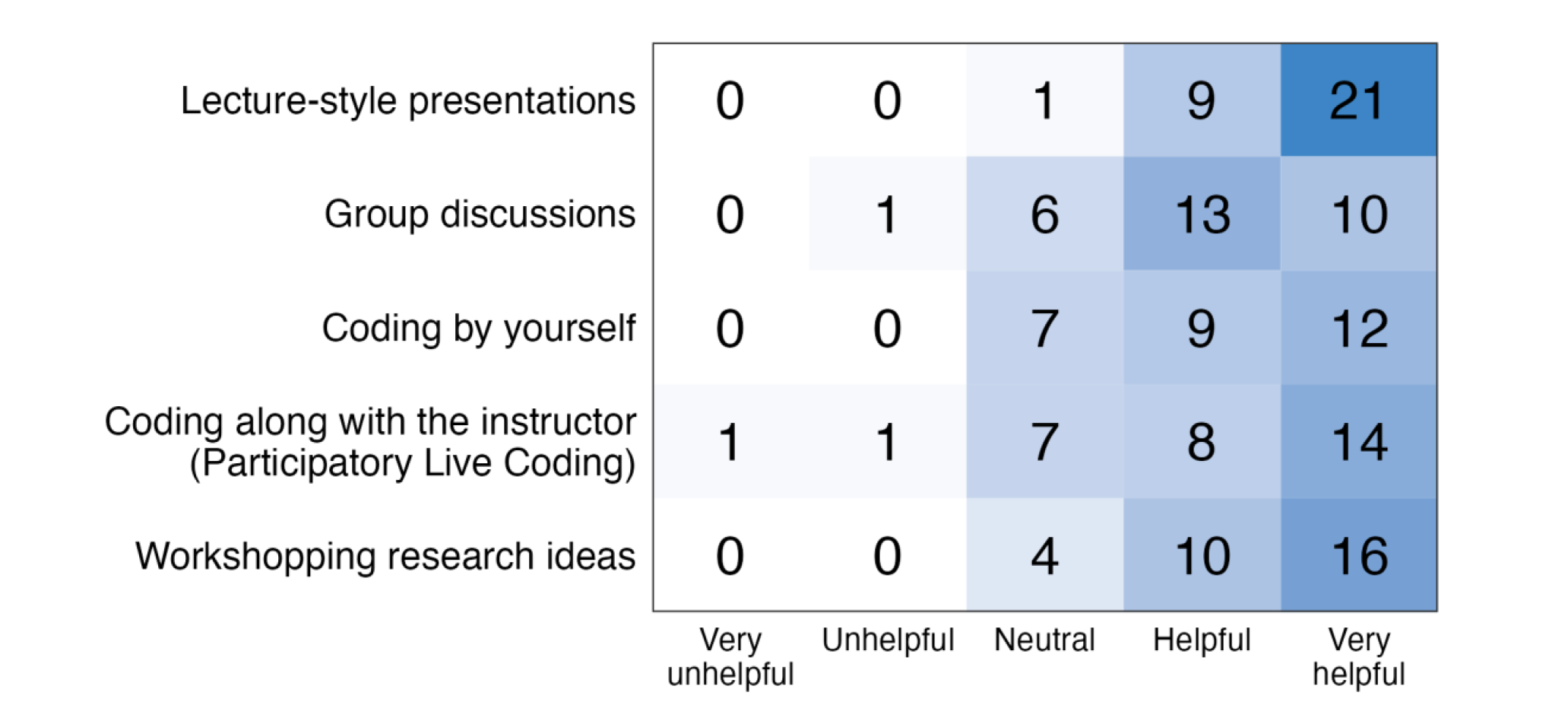
Participants’ ratings of the helpfulness of workshop activities. Overall, lectures were preferred over other activities (p=0.003, exact Wilcoxon signed rank test). Participatory live coding was considered helpful (mean response ∼4.06/5, p=0.015, exact binomial test), although 1 participant considered it unhelpful and 1 participant considered it very unhelpful.

**Figure 5.**
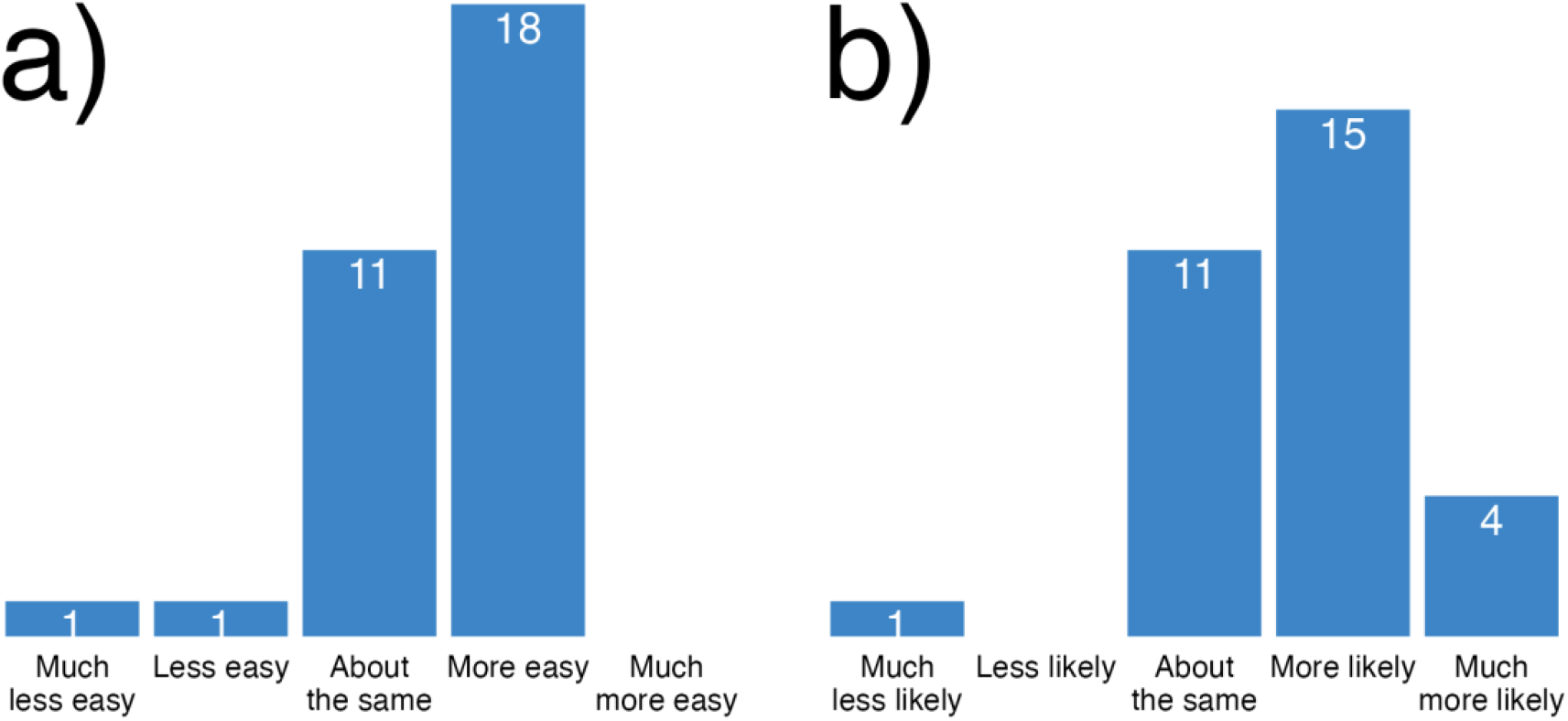
Participants’ perceptions of participatory live coding, and whether it will make implementing models easier (a) or make them more likely to engage (b). When asked specifically about participatory live coding, the null hypotheses of equal positive and non-positive outcomes could not be rejected; a non-significant number of participants indicated that this activity would make it easier to implement the models in the future (p=0.237, exact binomial test), or that this activity would make them more likely to engage with the models in the future (p=0.141, exact binomial test).

Participants’ feelings about participatory live coding were probably influenced by challenges during the administration of the workshop. When asked an open-ended question about workshop challenges, many participants indicated that it was difficult to engage with the live coding activity, citing the overall pacing of this activity as the primary contributing factor. This was true even amongst participants who found the activity helpful. One participant said, “[It was] very hard to follow the instructor while entering the code myself. I found [it] far more useful to just listen to the instructor and practice the code later.” Another stated, “I personally [am] the kind of learner who wants to walk step by step to understand why we’re making some decisions when following the tutorials. Because of this, when the pace was faster I found myself lost quite easily and sometimes I had to just disconnect from the instructor for a few minutes to read the tutorials on my own and type things on the command line in order to catch the main idea again.” If the duration of the workshop could be extended so that more time was allocated to the coding activities, participants may rate these activities more highly.

## Discussion

The value of computational workshops for researchers may seem self-evident. Researchers tend to like learning new things, and workshops that take place over the course of multiple days, or those that involve travel, require a substantial investment from participants and instructors. Individuals who do not expect to benefit from a workshop are unlikely to enroll, or be admitted after completing an application process. However, these factors do not necessarily imply that the specific goals of the workshop instructors or participants will be met during the workshop. It is entirely possible that a mismatch between the goals of instructors and participants might lead to some goals not being met. Additionally, workshops may be ineffective or even detrimental if they are perceived to be boring or too difficult. To learn more about the experiences of participants and improve workshops, instructors often administer surveys. However, in absence of a common set of goals for computational workshops, it is difficult to compare or assess the value of particular instructional techniques across different workshops.

Here, we focus on one key element of a successful workshop: its ability to motivate participants to use the methods and tools presented during the workshop. We note that even if a workshop is beneficial to participants in some ways, it may not motivate participants to engage with specific methods or improve their likelihood of doing future work in the field. It may even be the case that all participants already intend to do such research, and their research trajectory is not influenced by the workshop at all. We sought to assess these factors with our surveys.

We were initially concerned that despite anonymization, participants may be reluctant to deliver negative feedback to workshop instructors who may be colleagues, peers, collaborators, or even friends. However, this did not appear to be the case for our survey responses; respondents freely shared their difficulties and frustrations with live coding, suggesting that positive assessments in other areas were likely honest.

In our analysis of the phylogenetic biogeography workshop, we found that participant and instructor goals were met, and participants considered themselves significantly more likely to engage with phylogenetic models of historical biogeography after having completed the workshop. These results suggest that the workshop was effective overall, pointing to the value of computational workshops for motivating researchers to use particular methods and tools in the field of evolutionary biology. However, determining whether this self-reported increase in planned engagement persists over time, or whether it ultimately results in associated research, will require a longitudinal study.

One essential finding from our survey was that participants were very interested in learning software tools, which was not advertised as a goal of the workshop during workshop recruitment (Fig. 1). While the workshop required learning about software tools, and participant goals were ultimately met, it is important to acknowledge that mismatches of this kind can happen, and instructors will not know about them unless they ask. It should not be assumed that participant and instructor goals will align, even with a descriptive advertisement or application process. Surveying participants prior to administering workshops should be commonplace. Some workshops already do this (e.g. the RevBayes workshops, Barido-Sottani et al., 2022), although the results of the surveys and the corresponding adjustments made by instructors are rarely published.

Our survey also indicated that some activities had a more positive impact on student perceptions and plans than other activities. According to participants’ self-reports, lecture-style presentations were the most helpful activity (Fig. 4). This is not surprising; research on classroom instruction suggests that students may *feel* like they learn more from lectures, whether or not they actually do (Deslauriers et al., 2019). Still, many participants highlighted the importance of learning about the benefits and drawbacks of different models, and cited this knowledge as one of the main reasons why they are more likely to use the models in the future. Lectures were essential for delivering this information.

The participatory live coding activity was also rated as helpful or very helpful by most participants, although it was considered less helpful than lectures (Fig. 4). However, we were not able to identify a statistically significant effect on motivation. When participants were asked if the activity would make it easier to implement the models in the future, or if the activity would make them more likely to engage with the models, the null hypotheses of equal positive and non-positive responses could not be rejected (Fig. 5). This may be due to a true non-effect, or simply a small sample size. It is also important to keep in mind that the participatory live coding activity may have other benefits in the researchers’ future endeavors apart from motivation, like providing them experience to draw from when performing coding tasks.

A small number of participants found participatory live coding unhelpful, thought the activity would make it harder to implement models in the future, or considered themselves less likely to engage with the models due to the activity. While it is rarely possible to tailor instruction to all participants simultaneously, these individuals were not the only ones to express concerns about the activity. Participants often referred to the pacing of participatory live coding when asked about workshop challenges. It is possible that a slower version of the activity may be more effective in improving confidence and encouraging participants to use biogeographic models in their future research. We urge other workshop instructors to investigate the effectiveness of participatory live coding and other instructional activities during their workshops, examining how different implementations may impact motivation.

Further studies should also address factors impacting participant experiences such as differing careers stages or knowledge levels, variability in workshop structure and content, and potential discipline-specific effects. A large-scale analysis across multiple workshops from different disciplines would strengthen our understanding of the value of computational workshops and participatory live coding more generally. Longitudinal studies should also be conducted to assess workshop participants’ perceptions and plans during the months or years after a workshop, and whether these plans actually result in the generation of new research.

These types of advancements would rely on a common set of workshop goals and standard tools to measure them. Here, we identified one essential element of a successful workshop: its ability to increase participants’ motivation to use the methods and tools presented during the workshop. We then measured this for the Phylogenetic Biogeography Workshop using pre- and post-workshop surveys. However, in the future, a common set of workshop goals should be constructed collaboratively by workshop stakeholders through focus groups and surveys, and validated survey tools specific to advanced-level workshops for researchers should be developed.

## Conclusions

In this study, we identify a key ingredient of a successful computational workshop for researchers: its ability to increase participants’ motivation to use the methods and tools presented during the workshops. Through surveying participants, we found that the Phylogenetic Biogeography Workshop encouraged participants to conduct biogeographic research using the tools and methods taught during the workshop, and participants felt more confident about several key topics, including biogeographic models and the software RevBayes. In terms of specific activities, conventional lectures were preferred by participants over coding activities, despite a prevalent interest in learning software tools. Participatory live coding was considered helpful or very helpful by most participants, but survey questions connecting participatory live coding to participant motivation did not show statistical significance. Our findings indicate that the Phylogenetic Biogeography Workshop was successful and provide insights into participants’ goals, perceptions, and plans. Although we present one case study based on a single set of workshops, we also highlight the need for future work in this area. Before we are able to expand the scope of pedagogical research in the workshop setting, workshop stakeholders should collaborate to clearly define a set of common workshop goals and develop standardized, validated tools for assessing the efficacy of computational workshops for researchers in the field of evolutionary biology.

## Additional

### Funding

We would like to thank the NSF for funding these workshops under grant NSF DEB 2040347 awarded to Michael Landis. This study was supported by the NSF through the Graduate Research Fellowship Program awarded to Sarah Swiston (NSF DGE 2139839). Any opinions, findings, and conclusions or recommendations expressed in this material are those of the author(s) and do not necessarily reflect the views of the National Science Foundation. The authors also wish to acknowledge the Siteman Cancer Center’s National Cancer Institute (NCI) Cancer Center Support Grant P30 CA091842, the Washington University Institute of Clinical and Translational Sciences Grant UL1 TR002345 from the National Center for Advancing Translational Sciences (NCATS), and the Institute for Informatics, Data Science & Biostatistics REDCap Support teams for supporting the Washington University instance of REDCap (Research Electronic Data Capture). NCI and NCATS are part of the National Institutes of Health (NIH).

## Acknowledgements

We would like to acknowledge the co-organizers of the phylogenetic biogeography workshops Felipe Zapata, Isaac Lichter-Marck, and Fábio Mendes, co-instructors Mike May and Ixchel González-Ramírez, and all of the workshop participants who helped make this study possible.

## Supplementary Material

Supplementary materials including surveys, recruitment tools, and analysis code are available from the Dryad Digital Repository: INSERT DRYAD LINK HERE. Materials are also available on GitHub: https://github.com/sswiston/workshop_survey. Raw survey results are excluded for the privacy of participants.

## Supplementary Materials S1: Pre-Workshop Survey

1. What is your current career stage? [multiple choice -- pick one]
  - Undergraduate student
  - Master’s student
  - Ph.D. student
  - Postdoctoral researcher
  - Faculty
  - Non-faculty professional
  - Other (specify)
2. What were your initial goals when you applied for this workshop? [checklist -- check all that apply]
  - Learn about the kinds of questions that phylogenetic methods of historical biogeography can answer
  - Learn more about different types of models
  - Learn more about software tools
  - Try out coding a model with the help of workshop facilitators
  - Decide whether to do future research using phylogenetic methods of historical biogeography
  - Discuss research ideas with workshop facilitators and other participants
  - Network with other researchers
  - Add the workshop to a resume or CV
  - I did not have any goals when I applied for this workshop
  - Other (specify)
3. Please rate your experience with the following topics and tools: [options: unfamiliar, somewhat familiar, competent, highly competent, expert]
  a. Probability and statistics
  b. Molecular phylogenetics
  c. Divergence time estimation
  d. Biogeographic models
  e. Bayesian models
  f. Markov chain Monte Carlo
  g. Unix command line
  h. RevBayes
4. During your time as a researcher, how often have the following events occurred? [options: never, rarely, sometimes, often, very often]
  - I wanted to use a research method, but stopped because the <u>software</u> was too difficult to use.
  - I used a research method even though the <u>software</u> was difficult to use.
  - I wanted to use a research method, but stopped because the <u>programming</u> was too difficult.
  - I used a research method even though the <u>programming</u> was difficult.
  - I wanted to use a research method, but stopped because the <u>statistical analysis</u> was too difficult.
  - I used a research method even though the <u>statistical analysis</u> was difficult.
  - I wanted to use a research method, but stopped because the <u>model</u> was too difficult to understand.
  - I used a research method even though the <u>model</u> was difficult to understand.
5. Do you plan to use phylogenetic methods of historical biogeography in your own research [yes / no / unsure]?
  a. If so, when do you plan to start using them? [options: I already use these models, within the next 3 months, between 3 and 6 months, between 6 months and 1 year, after 1 year]
  b. How do you plan to apply these methods in your own research? [open-ended]

## Supplementary Materials S2: Post-Workshop Survey

1. Did this workshop meet your expectations? Why or why not? [open-ended]
2. For each of the following topics and tools, please indicate how your level of confidence in your ability to use the tool has <u>changed</u> after engaging in the workshop: [options: much less confident, less confident, about the same, more confident, much more confident]
  a. Probability and statistics
  b. Molecular phylogenetics
  c. Divergence time estimation
  d. Biogeographic models
  e. Bayesian models
  f. Markov chain Monte Carlo
  g. Unix command line
  h. RevBayes
3. Of the tools or skills addressed in the workshop, what is one thing you could implement in your research relatively easily? [Open-ended]
4. How likely are you to use skills or tools that you <u>acquired during the workshop</u> in the future? [options: very unlikely, unlikely, neutral, likely, very likely] Why? [open-ended]
5. If you plan to use skills or tools acquired during the workshop, which one(s) do you plan to use? [open-ended]
6. Are you more or less likely to engage with phylogenetic models of historical biogeography <u>after having taken the workshop</u> (compared to how you felt before)? [options: much less likely, less likely, about the same, more likely, much more likely] Why? [open-ended]
7. How helpful were these specific activities to your learning? [options: very unhelpful, unhelpful, neutral, helpful, very helpful, N/A]
  a. Lecture-style presentations
  b. Group discussions
  c. Coding by yourself
  d. “Coding along” with the instructor
  e. Workshopping research ideas
8. Did you encounter any challenges during the workshop, and how did you handle them? [Open-ended] **The term “participatory live coding” refers to activities where the instructor was typing code that everyone could see, and you “coded along” on your own computer**.
9. Do you think the <u>participatory live coding</u> activity will make it easier to implement phylogenetic models of historical biogeography in the future? [options: much less easy, less easy, about the same, more easy, much more easy, I did not participate in this activity]
10. Do you think the <u>participatory live coding</u> activity made you more or less likely to engage with phylogenetic models of historical biogeography? [options: much less likely, less likely, about the same, more likely, much more likely, I did not participate in this activity]

## Supplementary Materials S3: Recruitment Emails

### Before Workshop

Hello workshop participants! My name is Sarah Swiston, and I will be one of the instructors for the upcoming workshop on phylogenetic methods for historical biogeography. As part of my dissertation, I’m doing a research project about the design and effectiveness of computational workshops. Here, I am attaching a pre-workshop survey. It asks briefly about your expectations for the workshop and previous experiences with various research topics and tools -- similar to the application, but designed to answer specific pedagogical research questions. The survey is completely optional and responses will be anonymized before analysis. It should take about five minutes to complete. Filling it out will help us design future workshops and contribute to general knowledge about the effectiveness of workshops in evolutionary biology. You are welcome to fill it out now, and it will stay open until the end of the first workshop coffee hour. There will also be a post-workshop survey, which I will announce at the end of the workshop. Feel free to ask me any questions you might have about this research project! Thanks!

### After Workshop

Hello workshop participants! I mentioned at the beginning of the workshop that, as part of my dissertation, I’m doing a research project about the design and effectiveness of quantitative workshops. I am now attaching a post-workshop survey asking how you feel about the workshop, topics, and activities. Once again, the survey is entirely optional, and responses will be anonymized before analysis. It should take about ten minutes to complete. Filling it out will help us design future workshops and contribute to general knowledge about the effectiveness of workshops in evolutionary biology. Even if you didn’t fill out the pre-workshop survey, it would still be helpful to complete this one. You are welcome to fill it out now, and it will stay open for a week in case you would rather fill it out on your own time. Feel free to ask me any questions you might have about this research project! Thanks!

## Supplementary Materials S4: Verbal Announcements

### Before Workshop

As part of my dissertation, I’m doing a research project about the design and effectiveness of computational workshops. Some of you might have noticed that we sent out a pre-workshop survey. It asks briefly about your expectations for the workshop and previous experiences with various research topics and tools -- similar to the application, but designed to answer specific pedagogical research questions. The survey is completely optional and responses will be anonymized before analysis. It should take about five minutes to complete. Filling it out will help us design future workshops and contribute to general knowledge about the effectiveness of workshops in evolutionary biology. You are welcome to fill it out now, and it will stay open until the end of the coffee hour. There will also be a post-workshop survey, which I will announce at the end of the workshop. Feel free to ask me any questions you might have about this research project! Thanks!

### After Workshop

Hello! I mentioned at the beginning of the workshop that, as part of my dissertation, I’m doing a research project about the design and effectiveness of quantitative workshops. I have now sent out a post-workshop survey asking how you feel about the workshop, topics, and activities. Once again, the survey is entirely optional, and responses will be anonymized before analysis. It should take about ten minutes to complete. Filling it out will help us design future workshops and contribute to general knowledge about the effectiveness of workshops in evolutionary biology. Even if you didn’t fill out the pre-workshop survey, it would still be helpful to complete this one. You are welcome to fill it out now, and it will stay open for a week in case you would rather fill it out on your own time. Feel free to ask me any questions you might have about this research project! Thanks!

